# Delayed dichromatism in waterfowl as a convenient tool for assessing vital rates

**DOI:** 10.1101/2024.06.04.597326

**Authors:** Adrien Tableau, Iain Henderson, Sébastien Reeber, Matthieu Guillemain, Jean-François Maillard, Alain Caizergues

## Abstract

Monitoring the number of individuals is by far the most popular strategy for studying the environmental factors that determine population dynamics and for measuring the effectiveness of management actions aimed at population recovery, control or eradication. Unfortunately, population size monitoring is inefficient in identifying the mechanisms underlying demographic processes and, in particular, in assessing the extent to which population growth rate is influenced by changes in adult survival rather than variations in reproductive parameters. In many waterfowl species, sexual dichromatism is observed in adults, while immatures of both sexes display a plumage pattern similar to that of adult females. In these species, the apparent proportion of males increases as the female-like immature males gradually take on the plumage of adult males. The difference between the apparent sex ratio before and after the young reach sexual maturity then provides information about the age ratio of a population. Using winter counts that distinguished between female-like and male-like individuals of two non-native populations of Ruddy duck *Oxyura jamaicensis*, a species that exhibits such a plumage pattern, we present a non-invasive method based on the apparent sex ratio to split population growth rate into adult survival and recruitment rates (the latter also referred to as productivity). This method can correctly detect annual changes in vital rates, supporting the assumption that counts conducted in an appropriate time window reflect the age structure of a population. We exemplify how the respective contributions of survival and productivity to the population growth rate are essential for understanding the processes behind demographic dynamics. Finally, we point out some best practices to correctly apply the “apparent sex ratio” method described here.

## 1 Introduction

Assessing the growth rate of natural populations is a first step towards a better understanding of the factors underlying their dynamics (e.g. Niel & Lebreton, 2005). It is also crucial for measuring the effectiveness of management actions that are implemented to restore, control, or eradicate populations (Shea & NCEAS Working Group on Population Management, 1998). Among the approaches available to managers to achieve these goals, those that rely on monitoring the number of individuals (counts) are by far the most popular (Rintala et al., 2022). In many cases, these methods allow the investigation of environmental factors underlying changes in population size, and thus help in the implementation of mitigating actions (Faillettaz et al., 2019). Unfortunately, relying solely on the monitoring of population abundance generally hinders the fundamental understanding of the demographic mechanisms underlying changes in population growth rates. More specifically, on the basis of counts alone, it is generally impossible to assess the extent to which population growth rate is affected by changes in adult survival rather than to variations in the production of young (Austin et al., 2000). To assess the relative sensitivity of population growth rate to factors affecting adult survival or reproductive parameters, demographers usually rely on the monitoring of individuals by capture-mark-recapture (Lebreton et al., 1992). In practice, however, capture and release of individuals cannot always be relied upon, as the legal status of the species precludes any additional disturbance (e.g. critically endangered species) or prohibits the release of live individuals (e.g. invasive species or pests). Even though capture-mark-recapture methods are efficient for assessing demographic parameters, they also have potential drawbacks, such as their invasiveness, which can affect behavior and thus survival or reproductive success, and their affordability when time and money are scarce. Genetic monitoring is a non-invasive alternative to capture-mark-recapture, but it is costly and requires large samples (Caniglia et al., 2011).

Most of the time, therefore, managers make the best of a bad job by using counts as a viable option for tracking changes in population size and then population growth rate to assess the relevance of management actions. Distinguishing between immatures and adults in counts allows estimation of adult survival rate, i.e. the proportion of breeders that have survived for one year, and productivity/recruitment rate, i.e. the number of immatures produced per breeder that reach sexual maturity, which are, by definition, recruited in the breeding population (Smith et al., 2001). This additional effort makes it possible to assess the relative influence of each of these components on population growth rate. However, only few species, at least in waterfowl, exhibit morphological differences between age groups that can be recognized from a distance, and then recorded during counts. Alternatively, assessment of age structure in hunting bags has been used to infer the role of decreasing reproductive success in population declines in a number of game species, including ducks and geese, but this suffers from intractable biases (Fox & Cristensen, 2018). Clearly, the latter approach is also not appropriate for protected/endangered species. This emphasizes the need to develop non-invasive alternatives that can correctly track vital rates.

Here we test the hypothesis that delayed sexual maturity in dimorphic waterfowl species is a reliable source of indirect information on the age structure of a population, allowing the tracking of changes in vital rates over time. To this end, we first implemented a model based on winter counts that distinguishes between male-like and female-like individuals to infer these vital rates. We then compared the outputs with other sources of information on vital rates to check that the estimates coming from the tested method are not biased and accurately reflect changes in vital rates over time. We used two non-native European populations of Ruddy duck (*Oxyura jamaicensis*) as a study model. Like many other duck species, Ruddy duck is dimorphic: newborn males look like females until the prenuptial molt, which takes place during late winter. As a result of the late prenuptial molt of immatures, the apparent proportion of males increases over the course of the wintering season. Such changes in the apparent proportion of males during this period are therefore directly related to the proportion of immatures in the populations and thus to the reproductive success of the previous breeding season. The “apparent sex ratio” method developed in this study was used to estimate adult survival and recruitment rates and to evaluate the effects of two different eradication strategies used in Great Britain and France, respectively.

## 2 Materials & methods

The Ruddy duck is a stiff-tailed duck native to the American continent. Starting from seven individuals acclimatized at the Slimbridge Wetland Centre in the 1940s (Gutiérrez-Expósito et al., 2020), a feral population began to establish in Great Britain and the first attempts at reproduction in the wild were observed in the 1960s (Figure 1). This feral population grew rapidly and spread across the country until it reached more than 5,000 individuals in the early 2000s. In the late 1980s, a feral population also began to establish on the continent, particularly in France, supposedly due to the arrival of British-born individuals (Muñoz-Fuentes et al., 2007). However, in contrast to observations in Great Britain, Ruddy ducks have not spread very much in France. Most observations and breeding attempts have been concentrated in the north-west of the country in ponds with a rich vegetation. Ruddy ducks commonly gather in large open waters to overwinter. In line with this, almost no Ruddy ducks were observed outside the lake of Grand Lieu (47.09°N, 1.67°W) during winter in France, which greatly facilitated the monitoring of this population. Ruddy ducks are considered a major threat to the endangered native White-headed duck (*Oxyura leucocephala*) in the south-western Mediterranean, as they hybridize and thus pose an increased risk of genetic pollution and genetic assimilation of the latter by the former (Muñoz-Fuentes et al., 2007). To reduce the risk of genetic pollution of the White-headed duck by the Ruddy duck, eradication measures were implemented in both Great Britain and France in the late 1990s (Gutiérrez-Expósito et al., 2020), followed by a European Ruddy duck eradication plan in 1999 (Hughes et al., 1999) (Figure 1).

**Figure 1:**
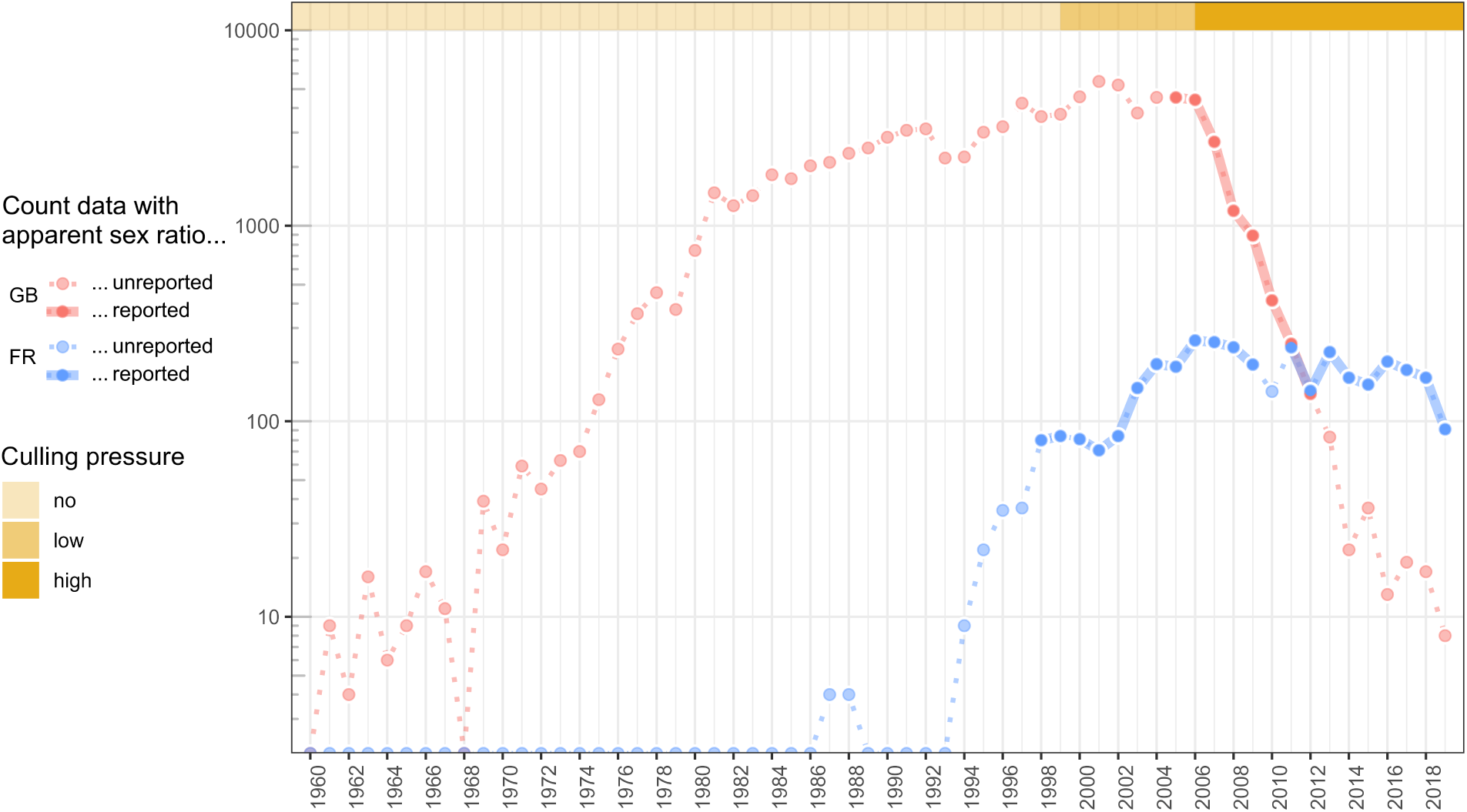
Temporal changes in the number of Ruddy ducks (logarithmic scale) in Great Britain (GB) and France (FR) from 1960 to 2019, with references to periods when apparent sex ratios were reported; data were collected exclusively during winter

The trends and status of the White-headed duck and Ruddy duck populations used to be assessed exclusively by counts. It was therefore difficult to assess the factors impairing the recovery of the former and the effectiveness of the eradication programme for the latter. In particular, counts alone cannot be used to assess the relative impact of changes in adult survival and recruitment rates on population growth rates, which is a prerequisite for identifying limiting factors.

Like many other duck species, both White-headed duck and Ruddy duck exhibit delayed dichromatism, i.e. young males acquire the colorful plumage typical of their species and can therefore be distinguished from females during the pre-breeding period, typically from mid-winter in the earliest individuals (Baldassarre, 2014). Delayed dichromatism generally explains the discrepancies in estimates of apparent proportions of males counted during winter compared to proportions of males counted during spring or males identified from individuals culled during winter (Figures 2 & 3).

**Figure 2:**
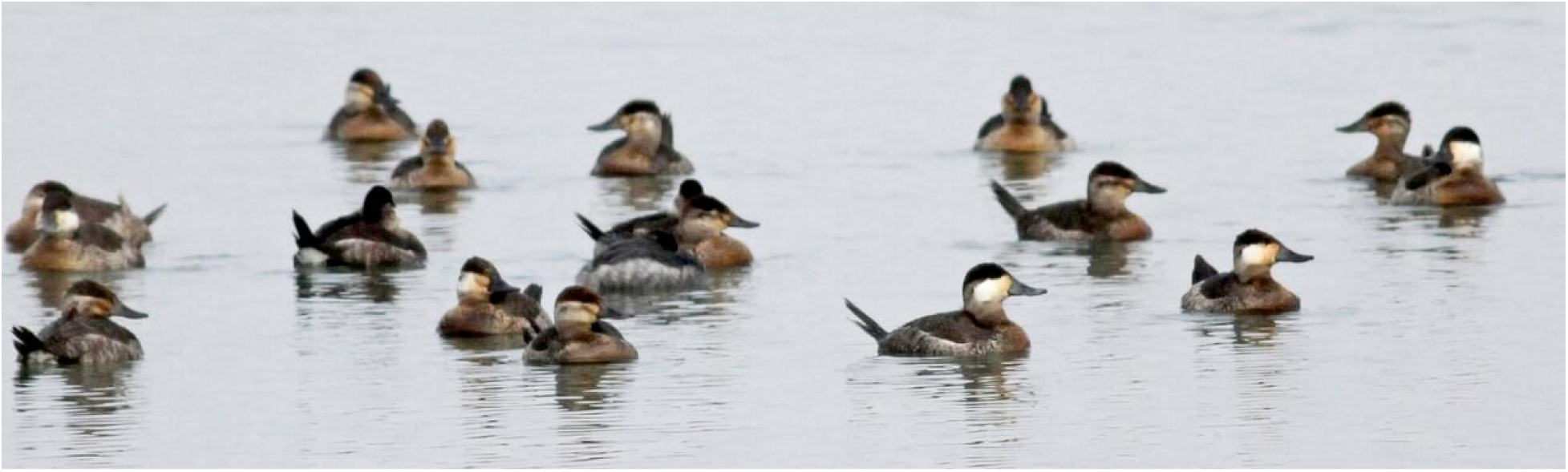
A flock of Ruddy ducks observed during winter, including ten female-like individuals with whitish striped cheek (some of which are immature males), four male-like individuals with white cheek and black cap (all adult males), three unidentified individuals (1^st^, 4^th^, and 10^th^ from left) © Jay McGowan - 3 February 2013 - Tompkins, New York, United States

Interestingly, delayed dichromatism allows the estimation of adult survival and recruitment/productivity by monitoring abundances and the seasonal switch of the apparent sex ratio before and after the molt of immature individuals.

### 2.1 Data formatting

The method requires the specification of three observable variables (see “Observations” in Figure 4). First, the size of Ruddy duck populations was monitored in both Great Britain (GB) and France (FR) by conducting extensive counts on the wintering grounds between 1 December and 31 January from 1960 to 2019. During this period, the individuals in the wintering grounds are highly detectable as they gather in open water. At least one count was carried out annually around mid-January. In some years, additional counts have also been carried out in December in GB and FR. Although the counted numbers are generally similar (variation coefficient of 10%), the peak counts are generally observed in January, suggesting that a small part of the population arrives late at the wintering grounds in some years. This conclusion prevents the estimation of detection probability using an N-mixture model (violation of the “closed population” assumption, see Costa et al. (2021)) and led the authors to retain the maximum count value as an estimate of population size (Figure 1).

**Figure 3:**
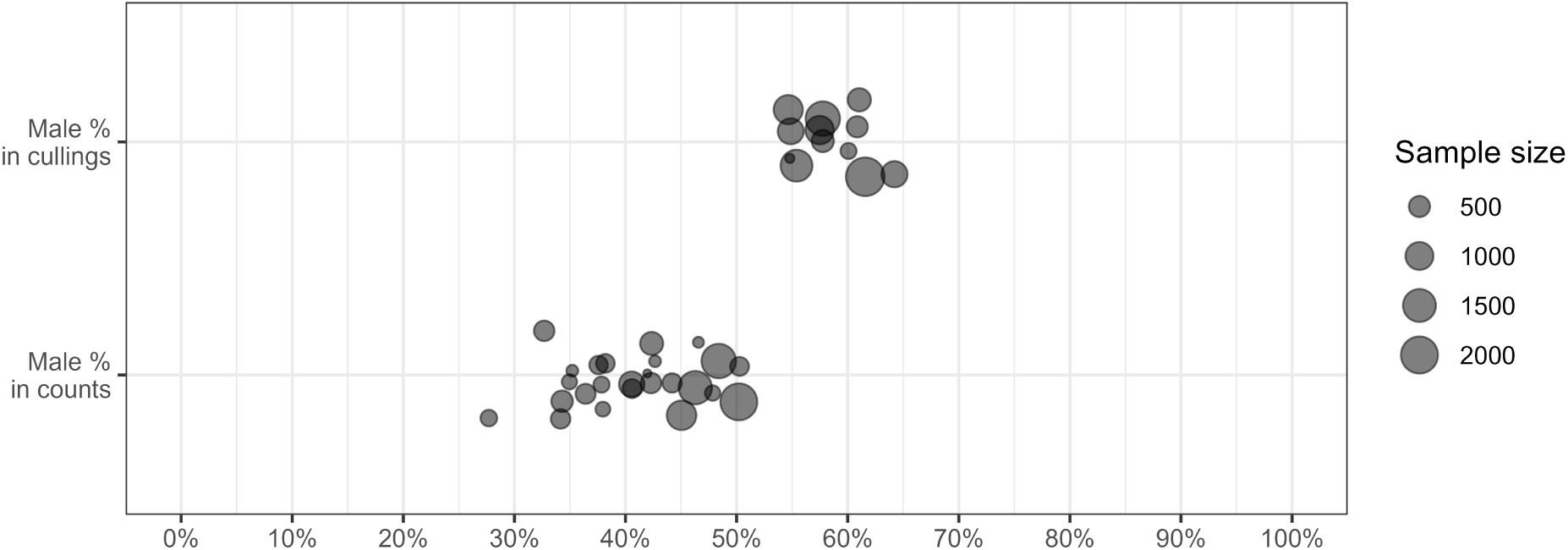
Proportions of males estimated by internal examination of individuals culled in a given year under the eradication programmes (Male % in cullings), and proportions of individuals showing male-like plumage counted during winter (Male % in counts); data from Great Britain and France are pooled together; discrepancies between the two estimates are due to delayed dichromatism (immature males looking like females before molting); we used such discrepancy to estimate survival and productivity/recruitment

**Figure 4:**
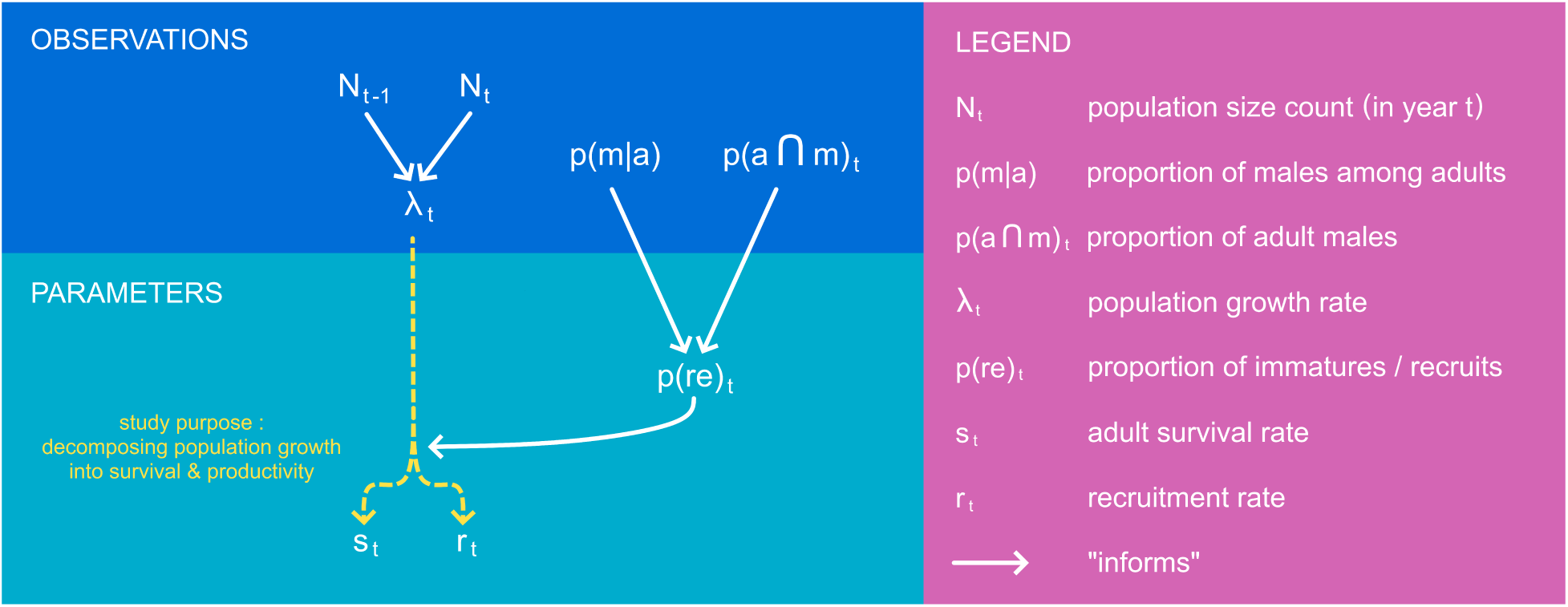
Overview of the “apparent sex ratio” method used to disentangle the population growth into survival (adult survival rate) and productivity (recruitment rate)

Second, the sex ratio in adults (alternatively, the proportion of males among adults), which is presumed to be stable over several years, is theoretically observable during spring when all males are sexually mature and display their colorful plumage. However, the poor detectability of this species at this time of year prevented from using such an approach. This variable was then determined directly on culled individuals, which can be aged and sexed in the hand with certainty. The absence of the bursa of Fabricius was used as a reliable criterion for determining an adult, and the presence of a penis indicates a male individual. In France, it was not possible to obtain sufficiently reliable estimates of sex ratio in adults due to the small sample size. In Great Britain, a preliminary analysis indicated no seasonal variation of the sex ratio in adults. Data from culled adults collected throughout the whole biological cycle were then pooled to assess annual sex ratios. There was also no statistical difference in adult sex ratio between years in which more than 500 adults were culled. This is consistent with previous findings showing that sex ratios in adult ducks tend to be very stable in the short term (Wood et al., 2021), but can fluctuate in the long term. Therefore, we pooled the data from all adults culled in the control programme to estimate the sex ratio in adults.

Third, the apparent sex ratio when immature males look like females was assessed annually by pooling all winter counts that distinguished between female-like and male-like individuals. It is assumed that this apparent sex ratio directly reflects the proportion of adult males in the population. Years with winter counts distinguishing apparent sex are a subset of the extensive count time series described previously (Figure 1). These counts were carried out in Great Britain from 2006 to 2012, which corresponded to a period of sharp population decline. In France, counts distinguishing apparent sex were carried out in 1999, 2001-2009, and 2012-2019. The French population grew rapidly in the first years of monitoring and then stabilized from 2006 onwards as a result of high culling pressure.

In both countries, culling was carried out under strict official control, so the exact number of individuals culled was known. Although the age ratio of birds culled before 2009 could not be determined with certainty in France, it was still possible to roughly categorize the culling pressure in both countries into three categories (Figure 1): “no culling” before 1999, as the culling rate on adults was mostly zero and always below 10% in both countries, “low culling” between 1999 and 2005, as the adult culling rate fluctuated around 20%, and “high culling” from 2006 onwards as the adult culling rate fluctuated around 50% (see Figure 8 in the Appendix section).

### 2.2 Inferring adult survival and recruitment rates from changes in population size and proportion of immatures

The method introduced in this study is called the “apparent sex ratio” method and is performed as a single model structured in two levels (see “Parameters” in Figure 4).

First, the population growth rate, which is the ratio of the population size over two consecutive years, must be determined. The method is also valid if only abundance indices are available. In parallel, the originality of the method consists in inferring the immature/recruit proportion of a population by analyzing the difference between the sex ratio among adults (which is stable over time), and the apparent sex ratio observed in counts carried out during winter (see Equations (1) & (2)). These counts are pooled according to the additive property of the binomial distribution.

Second, combining the population growth rate with the proportion of immatures yields estimates of adult survival and recruitment rates, which are latent variables, see Equation (3) and Figure 4. If the absolute value of the population size is known, the number of adults and the number of recruits can be estimated, see Equation (4).

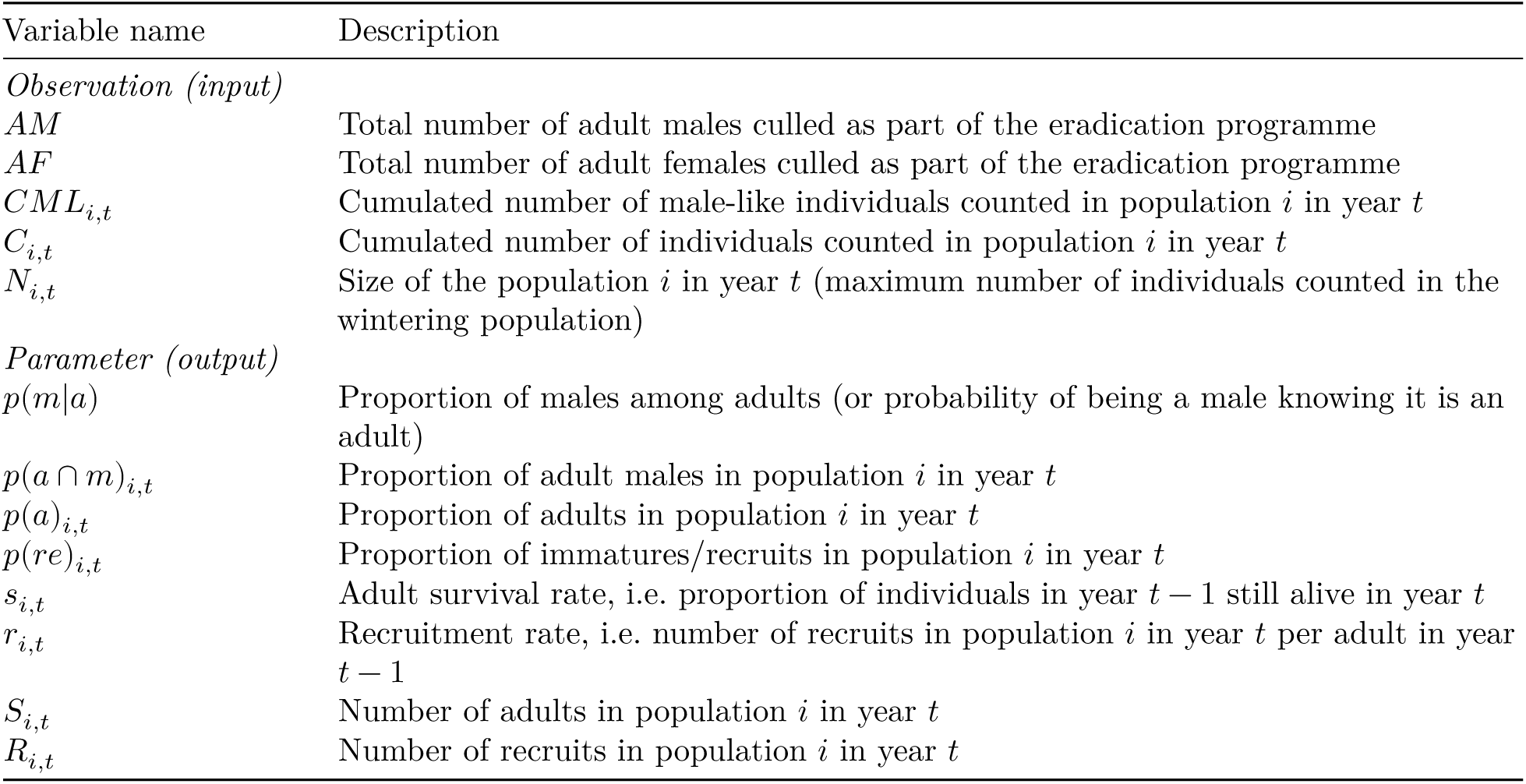

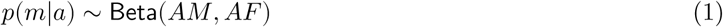

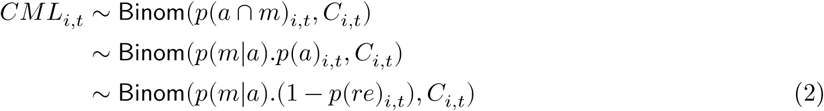

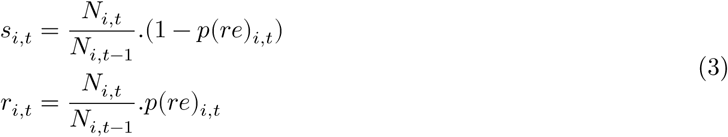

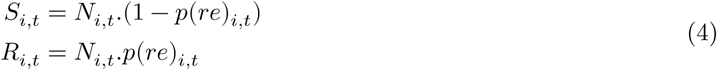

### 2.3 Validating the “apparent sex ratio” method

The method is based on three assumptions. The first assumption is that the population size is known exactly. As explained in the Data formatting Section, the detection probability of birds occupying wintering grounds is reasonably close to 100%, so detection issues should be negligible in the present case study. A negative bias in population size may nevertheless occur if not all birds arrived at the wintering grounds at the time of the exhaustive count survey. If the bias is constant over years, this is not an issue as the population growth rate is a relative quantity and can be estimated using abundance indices. However, if some birds arrive late only in some years, the population growth rate may be positively or negatively affected, depending on the year in which such a bias occurs. This assumption is discussed further, but remains specific to this case study. This bias can be avoided in other studies by conducting repeated counts around the peak of occupancy of wintering grounds. In the case of imperfect detection, true population size can be assessed using repeated counts within an N-mixture framework (see Costa et al., 2021 for details).

The second assumption is the stability of the adult sex ratio over time. Violation of this assumption would lead to an increasing bias in the vital rates with the drift of the adult sex ratio over time. An examination of the stability of the sex ratio over the time series available in GB confirms that this assumption of is valid (see Data formatting Section). The stability of the adult sex ratio was observed at least over a decade, which is in line with the literature (e.g. Wood et al., 2021).

The third and most important assumption is that the male-like individuals observed during winter correspond exclusively to adult males, as immature males are mistaken for females at this time of year. A violation of this assumption occurs when a significant proportion of immature males have acquired male plumage prior to the counts considering apparent sex. This violation would be detected through estimates of adult survival rate that are higher than expected for this species and, conversely, a recruitment rate biased low. A strong corruption of this assumption would even lead to adult survival estimates higher than one and negative recruitment rates for closed populations. A second violation occurs when adult males are less detectable than other individuals, but nothing indicates that such an issue would occur. This would be detected through an overestimation of recruitment rates and low or even negative adult survival rates.

The latter assumption was assessed by testing the likelihood of the estimates of both vital rates. For adult survival rate, we first checked whether the values were contained in the interval [0; 1], and compared our estimates with those in the literature (e.g. Buxton et al., 2004; Krementz et al., 1997; Nichols et al., 1992, 1997). Validation of the recruitment rate was more difficult because if the lower limit is 0, the upper limit is a product of all maximum values of its individual components, i.e. nesting rate, nesting success, clutch-size, hatching success, pre-fledging survival, and fledging survival. Although some of these reproductive parameters may be known for some duck species, the last one is never mentioned to our knowledge (e.g. Baldassarre, 2014).

To address this problem, we developed an indirect approach to estimate the maximum expected recruitment rate without exploitation and under the assumption that the level of population density has no influence on the components of the recruitment process. In practice, maximum recruitment rate was estimated from the difference between the maximum growth rate and the maximum survival rate of adults, see Equation (5). This relationship resulted from a simple consideration for a closed population: the population size in year 𝑡 is equal to the number of adults that survived the whole year 𝑡 − 1 plus the offspring produced in year 𝑡 − 1 that survived to the reproductive period of year 𝑡, i.e. recruitment in year 𝑡 (Flint, 2015). In an open population, adult survival and recruitment rates are confounded with adult and recruit migration rates, respectively, but this does not change the equation. This relationship becomes more complex when a species with delayed maturity is considered, see Robertson (2008).

| Variable name | Description |
| --- | --- |
| $N_t$ | Number of adults in year $t$ |
| $D_t$ | Number of adults dead during year $t$ |
| $R_t$ | Number of recruits in year $t$ |
| $s_t$ | Adult survival rate, i.e. proportion of individuals in year $t - 1$ still alive in year $t$ |
| $r_t$ | Recruitment rate, i.e. number of recruits in year $t$ produced per individual in year $t - 1$ |
| $\lambda_t$ | Growth rate of the population between year $t - 1$ and year $t$ |

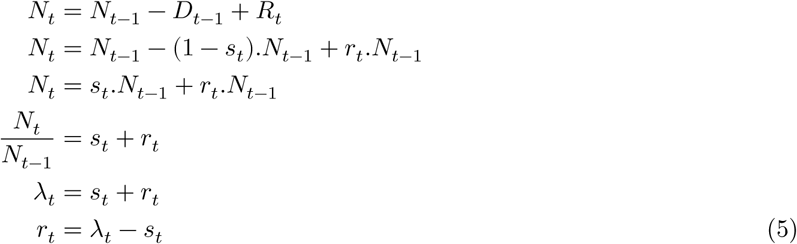

The maximum survival rate of adults was defined as a uniform distribution on the interval [0.7; 1]. The lower limit corresponded to the upper range of survival rates observed in waterfowl species of similar weight, and the upper limit was set to 1, as survival rates of long-lived waterfowl species can be very high (Buxton et al., 2004; Koons et al., 2014; Krementz et al., 1997; Nichols et al., 1992, 1997).

For both populations, the maximum growth rates occurred during their geographic expansion phase, which preceded the beginning of the eradication programmes (Figure 1). To estimate robust maximum population growth rates for both populations, we smoothed the annual growth rates over a consistent time period by using a linear regression on the logarithm scale, see Equation (6). We discarded the data for Great Britain before 1972 as the estimates were likely noisy when the population was low (below 50 individuals) (Figure 1). After reaching the threshold of 1,000 individuals, population growth in Great Britain (GB) showed a strong inflexion, although culling had not yet started (Figure 1). This observation suggests that beyond 1,000 individuals, a negative density-dependent process probably took place and led us to consider only the first sequence of the time series to infer maximum growth rate in Great Britain, i.e. 1972-1981. For the French (FR) population, the sequence without culling pressure covered the period 1994-1999.

| Variable name | Class | Description |
| --- | --- | --- |
| $N0_{i,J}$ | Parameter | Intercept of the regression model |
| $\lambda_{i,J}$ | Parameter | Average population growth rate over a restricted time interval $J$ for a population $i$ (in $year^{-1}$) |
| $t \in J$ | Index | Year index within the time interval $J$ |
| $\sigma_{i,J}$ | Parameter | Standard deviation of the regression model |

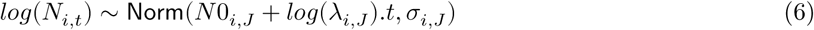

Estimating vital rates within an acceptable range, as defined above, would not prove that our modelling approach correctly reflects actual interannual variability. To determine this, we compared the proportions of immatures and the vital rates derived from counts with those derived from culling, i.e. from individuals culled as part of the eradication programme. As we restricted the dataset to years with more than 100 individuals culled during winter for the sake of precision, the analysis only covered five years of the GB population time series. The proportions of immatures were estimated from the culled individuals by checking the presence of the bursa of Fabricius, which is only present in immature individuals (Hochbaum, 1942); see Equation (7). Adult survival and recruitment rates were then derived from the proportion of immatures and the Equation (3).

| Variable name | Class | Description |
| --- | --- | --- |
| $SI_{i,t}$ | Observation | Number of immatures culled in the wintering population $i$ in year $t$ |
| $S_{i,t}$ | Observation | Number of individuals culled in the wintering population $i$ in year $t$ |

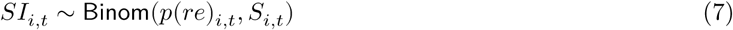

### 2.4 Assessing the effects of culling strategies

During the period when culling pressure was high (i.e. from 2006 onwards, see Figures 1 and 8), eradication strategies in Great Britain and France differed. In Great Britain, most individuals were culled during winter (53.4% of adults were culled before 30 May), whereas in France, most individuals were culled during the breeding season (81.6% of adults were culled after 30 May). We investigated whether these strategies had different effects on the populations by comparing the resulting growth rates and the relative contributions of both vital rates to them (by comparing the average values during the period of high culling pressure with proxies for the maximum vital rates estimated when both populations reached their maximum growth, see previous section). In France, a LIFE project was implemented from 2019 to intensify the culling pressure, especially during winter and spring. Therefore, we excluded the FR time series from 2019 onwards in order to compare homogeneous culling strategies.

### 2.5 Statistical framework

We used the Bayesian framework to implement the core model, which estimates vital rates using the “apparent sex ratio” method, and both validation models estimating maximum growth rates and vital rates from culled individuals. The Bayesian framework is both straightforward and efficient at propagating errors through the parameters. We used uninformative priors for all parameters. Since the maximum growth rate is a life history trait expected to be stable between populations of the same species, we used an uninformative hierarchical prior for this parameter. We generated three chains of length 500,000, with a thinning of 10 to avoid autocorrelation in the samples, and discarded the first 2,000 samples as burn-in. We checked the chain convergence using the Gelman and Rubin convergence diagnostic (R<1.1, Gelman & Rubin (1992)). The models were fitted using NIMBLE (de Valpine et al., 2017) run from R (R Core Team, 2022). The values **X[Y; Z]** reported in the Results section are the medians and the corresponding limits of the 95% credible interval of the posterior distributions.

## 3 Results

### 3.1 Estimating vital rates

Over the period 1960-2019, population size fluctuated between 0 and 5,473 in Great Britain, and between 0 and 259 in France. During this period, 12,316 Ruddy ducks were culled in GB, and 2,246 in FR. Of these, 8,440 adults were used to determine the male proportion among adults: 0.60 [0.59; 0.61]. In GB and FR, 12 and 55 independent count events considering apparent sex ratio were used to estimate vital rates over 7 and 18 years, respectively.

The “apparent sex ratio” method successfully provided estimates and associated uncertainties for the latent variables: proportion of immatures, and then, adult survival and recruitment rates (Figure 5).

**Figure 5:**
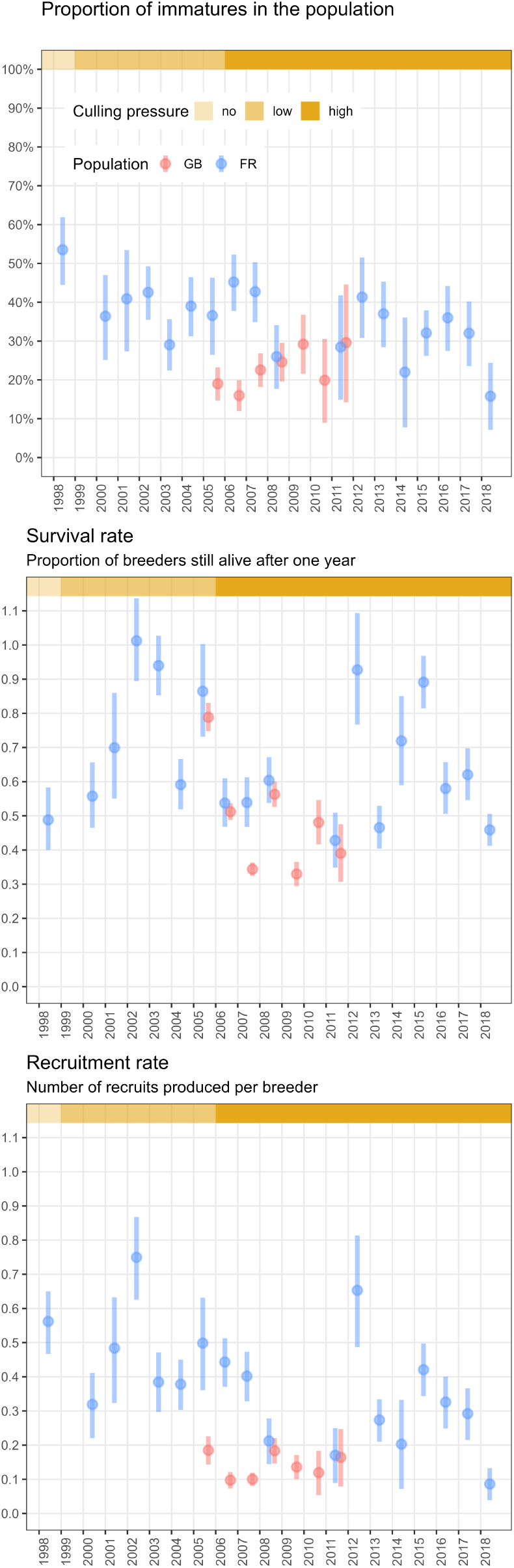
Temporal changes in the proportions of immatures and vital rates following culling pressure in the populations of Great Britain (GB) and France (FR); the vertical bars represent the 95% confidence intervals

The model estimated the proportion of immatures between 0.16 [0.07; 0.24] and 0.54 [0.44; 0.62], depending on the population and year. The lowest values were similar in both populations, but the range of the proportion of immatures in the GB population was much smaller (upper value GB: 0.30 [0.14; 0.45] & FR: 0.54 [0.44; 0.62]). The proportion of immatures in the GB population was stable over time, while a slight but significant decrease was observed in the FR population.

Adult survival rates ranged between 0.33 [0.29; 0.37] and 1.01 [0.89; 1.14]. None of the estimates were significantly outside the range of a survival rate defined without immigration [0; 1]. No trend in adult survival rate was observed in either population, but the patterns were different: adult survival in GB was lower and showed less interannual variability than in FR.

Recruitment rates ranged between 0.09 [0.04; 0.13] and 0.75 [0.63; 0.87]. All estimates were above 0, which is consistent with reality. Furthermore, the maximum recruitment rate was 0.68 [0.36; 0.78] (see 3rd Results section), and no estimate was significantly outside the range of recruitment rates defined without immigration [0; 0.78]. No trend was observed for the GB population over the seven years available, but the recruitment rate decreased for the FR population, although it was noisier than the proportion of immatures. As with survival rate, the GB population also showed lower recruitment rate values with less variability than the FR population.

Lower values and lower variability of both adult survival and recruitment rates estimated in the GB population than in the FR population probably explain the different trends of the two Ruddy duck populations: a sharp and constant decline in the GB population versus a slow and variable decline in the FR population (Figure 1). In the FR population, the range of recruitment rates (min/max difference: 0.66) was larger than the range of survival rates (min/max difference: 0.58). Conversely, the range of survival rates (min/max difference: 0.46) in the GB population was much larger than the range of recruitment rates (min/max difference: 0.08). This suggests that the variability in population growth rate in FR was mainly determined by recruitment, whereas in GB it was mainly determined by changes in adult survival.

### 3.2 Testing the reliability of the estimates

In GB, the proportions of immatures derived from the counts were positively correlated with those derived from the culled individuals (Figure 6). This result therefore supports the hypothesis that the proportion of female-like individuals in the overwintering populations is efficient for the assessment of age structure and thus productivity. The correlation was even stronger when looking at vital rates (Figure 6), but this was expected because the two methods for estimating these vital rates had a common component, namely the population growth rate (see Equation (3)). Interestingly, a 1:1 correlation was not achieved for any of the parameters. The proportion of immatures derived from the counts was always lower than the proportion derived from the culled individuals.

**Figure 6:**
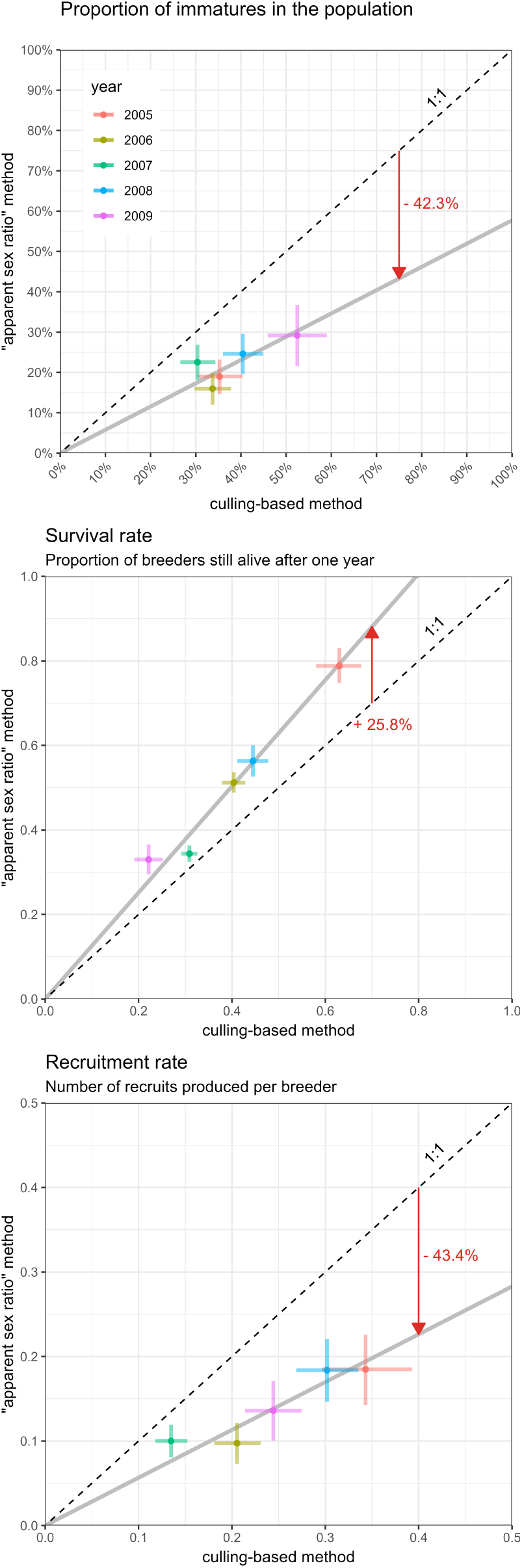
Relationship between parameter estimates obtained from counts (“apparent sex ratio” method) and those obtained from culled individuals (culling-based method); only five years were available for the GB population; the bars represent 95% confidence intervals and the red arrow indicates the direction and average difference between the two methods

### 3.3 Assessing the demographic response to culling strategies

When there was no culling pressure, maximum growth rates were very similar for the two populations (Figure 7), namely 1.45 [1.36; 1.55] and 1.52 [1.25; 1.88] for GB and FR, respectively. These values corresponded to population increases of 45% and 52% per year, respectively. Assuming a maximum adult survival rate of 0.85 [0.7; 1] for both populations, we derived maximum recruitment rates of 0.60 [0.42; 0.78] and 0.68 [0.36; 1.06] for GB and FR, respectively.

**Figure 7:**
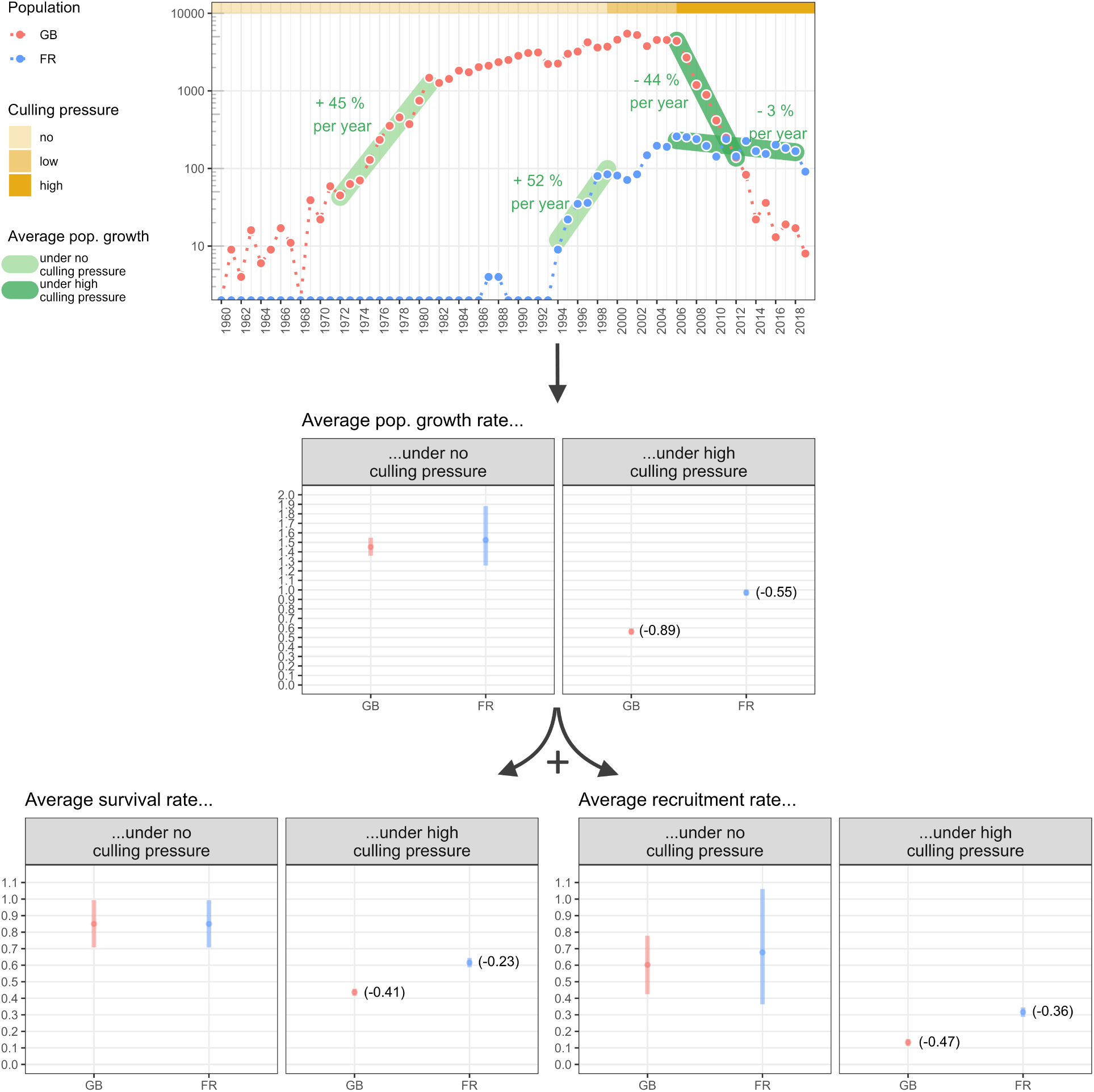
Average effects of culling pressure and different culling strategies (GB versus FR) on population growth rates (derived from counts only), and on adult survival and recruitment rates (derived from counts and reports of apparent sex ratios); the culling pressure for the GB population occurred mainly during the pre-breeding season, while for the FR population it was mainly concentrated during and after the breeding season; adult survival and recruitment rates under high culling pressure were averaged over a period in which consistent culling pressure was observed; the vertical bars show 95% confidence intervals

Despite the low number of sex- and age-determined culled individuals, the proxies of culling pressure showed significant changes over time (Figure 8). Therefore, we categorised culling pressure according to this intensity. High culling pressure occurred from 2006 onwards in both populations (although the signal was noisy). However, despite the similar level of culling pressure, the culling strategies of the two populations and their outcomes differed considerably (see 4^th^ Materials & Methods section). Under high culling pressure, the growth rate decreased to 0.56 [0.53; 0.59] and 0.97 [0.94; 1.00] for GB and FR, corresponding to a decrease of 44% per year and 3% per year, respectively. The GB population declined significantly during the high culling pressure, while the FR population was only stabilised.

Adult survival rates under high culling pressure reached 0.44 [0.42; 0.46] and 0.62 [0.59; 0.64] for GB and FR, respectively, and the recruitment rates decreased to 0.13 [0.11; 0.15] and 0.32 [0.29; 0.34], respectively. The resulting decrease in the growth rate of the GB population corresponded to a similar average decrease in adult survival rate (-0.41) and recruitment rate (-0.47), whereas the stabilisation of the FR population was due to a decrease in recruitment rate (-0.36) rather than a decrease in adult survival (-0.23). Overall, the culling effort in GB had a strong impact on both vital rates, leading to a greater population depletion than in FR.

## 4 Discussion

Disentangling the contribution of vital rates to population growth rate is a key step towards better understanding a population response to management actions (Williams et al., 2002). We exploited delayed sexual dichromatism to develop a new approach based on counts to disentangle the growth rate of a population into its two major structural components: adult survival and recruitment rates. The development of the “apparent sex ratio” method is based on a unique data set of two comparable populations of the same species, the Ruddy duck. Both populations were monitored in a similar way from their introduction to a period of intense culling pressure, but the culling strategies differed markedly. The very similar demographic trajectories and properties of both populations under similar ecological conditions made it possible to assess the response to both culling pressure and culling strategies.

### 4.1 Monitoring adult survival and recruitment rates

For both populations, vital rate estimates were within the same range, showing some consistency in the results obtained using the “apparent sex ratio” method. The greater variability observed for the FR population was expected as the time series covered a wider range of culling pressure than that of the GB population. We found no temporal correlation of demographic parameters between the two populations, suggesting that regional (e.g. weather) rather than large-scale factors (e.g. climate) were predominant. This is consistent with previous findings showing, for example, that in waterfowl both breeding success and juvenile survival depend on the onset of the laying period, which is closely linked to local weather (e.g. spring temperature, cumulative rainfall, water levels) (Blums & Clark, 2004; Dzus & Clark, 1998; Folliot et al., 2017) and local changes in predation pressure (Jaatinen et al., 2022).

The counts considering apparent sex used to implement our model took place in mid-winter, at a time when some immature males may be initiating their molt. This could partially violate the assumption that the male-like individuals were exclusively adult males, and thus potentially bias our estimates. However, as the survival estimates are in the range of those reported for similar species (Buderman et al., 2023; Buxton et al., 2004; Krementz et al., 1997; Nichols et al., 1997), this assumption appears to be reliable. Nevertheless, the time window for the counts must be chosen carefully and must match the time period before the immature males begin to molt.

Errors in the estimation of the population growth rate would also lead to absurd values of vital rates. This could typically happen if the extensive winter census used to estimate population abundance takes place before the arrival of the entire population in some years. In the present case study, the systematic count conducted in mid-January seemed to avoid this pitfall. When applying the method to other species, the approximate date on which all individuals reach their wintering grounds should be determined (e.g., from ringing or satellite-tagging) to ensure an unbiased estimate of population size.

In some years, the estimated values for adult survival were outside the range of expected values for a closed population, although they were never significantly above 1. One possible explanation for such discrepancies is that individuals from other populations moved to France in those years. Although ringing data suggest that Ruddy ducks in GB do not generally undertake long-distance seasonal migrations (Henderson, pers. comm.), we know that migration events have occurred in at least some years, as continental populations of Ruddy ducks in Europe are genetically closely related to the introduced GB population (Muñoz-Fuentes et al., 2006). Consistent with this, the outliers in survival and recruitment rates observed in FR in 2002 and 2012 suggest that immigration events may have occurred.

The theoretical maximum population growth rate is an inherent trait related to the demographic characteristics of a species (Dmitriew, 2010; Niel & Lebreton, 2005). This trait is expected to be similar in all Ruddy duck populations around the world. In our study, we observed that the maximum growth rates of the two populations were very close to each other, which was expected as both populations were exposed to similar ecological conditions, i.e. no harvest and similar breeding conditions. In addition, before the eradication programmes were initiated, the two populations had population growth rates much closer to the expected maximum values than any other native duck species in the same areas during the same period. Prior to the eradication programmes, the survival rates of both populations were in the upper range of those observed for other diving duck species (e.g. ∼ 0.8 for Common pochard (Folliot et al., 2020; Nichols et al., 1997)). Predation on diving duck nests was found to decrease over the course of the breeding season in a French fishpond complex (Bourdais et al., 2015). As Ruddy ducks generally breed later than Common pochards and Tufted ducks, both a higher survival rate and greater nesting success could be responsible for the high growth rates of Ruddy duck populations in the absence of culling. The “apparent sex ratio” method provided estimates of adult survival and recruitment rates that were not only consistent with observed increasing population sizes, but also compatible with both declining phases following the implementation of eradication programmes and observed differences in population growth rates under different culling strategies.

Assuming that culling data provide a good picture of interannual variability in age structure (Fox et al., 2014), the strong correlation between estimates derived from counts and estimates derived from culled individuals demonstrates the ability of the “apparent sex ratio” method presented here to properly capture interannual variation in demographic parameters such as survival and recruitment rates. The fact that the observed correlation was strong despite being based on a short time series strengthens its robustness. In addition, the strength of the relationship between the parameters obtained with the two approaches also indicates that important assumptions of our model were met, such as the constant adult sex ratio. The temporal autocorrelation of the adult sex ratio could be explained by the fact that it includes many age cohorts, which makes it structurally strong. In the long term, there may be substantial fluctuations in the adult sex ratio, but preliminary results suggest that it is not necessary to monitor and update it annually. The proportion of males among adults estimated in the present study (i.e., 0.60) is within the range of values observed in native Ruddy duck populations of North America (0.62 in Bellrose (1980)) and in other duck species (Wood et al., 2021). The “apparent sex ratio” method consistently yielded recruitment rates that were twice as low as the method based on culling data, a result consistent with higher vulnerability of immatures to shooting (Fox et al., 2014). In North America, for example, immature waterfowl were found to be 1.3 to 2.6 times more vulnerable to hunting than adults (Bellrose, 1980).

As no estimates of adult survival were available in the literature (see Buderman et al., 2023), we were unable to properly assess the accuracy of the “apparent sex ratio” method for estimating this parameter in Ruddy duck. Nevertheless, there is evidence that the method is not highly biased, as explained above. Thus, the “apparent sex ratio” method could be much more efficient for monitoring key demographic parameters than alternative methods, such as those based on culling/hunting bag data, which have been shown to be unreliable (Fox et al., 2014), even when focusing on the end of the hunting season to limit potential bias (Fox et al., 2016).

### 4.2 Evaluating eradication strategies

Both populations responded to culling pressure with a significant decline in their growth rate, meaning that the increased mortality caused by the culling was not compensated for by an increase in natural survival or improved breeding success. Culling was therefore effective in affecting the demographic trajectory of Ruddy duck in both cases. However, the decline in population growth was much steeper in GB, where the population declined steadily and sharply, than in FR, where the increase was halted and the population size remained more or less stable. The greater decline in population growth observed in GB compared to FR, was due to a greater decline in adult survival and recruitment rates. Overall, adult survival and recruitment rates appeared to be significantly affected by culling in both countries. However, culling pressure targeting future breeders (pre-breeding culling strategy applied in GB) appeared to be much more effective in reducing both survival and recruitment rates than culling pressure targeting breeders and newly born individuals during the breeding and rearing season (breeding culling strategy applied in FR). However, it cannot be excluded that population size also influenced the presumed effect of the culling strategy as the population size in GB was much larger than in FR.

As expected, the pre-breeding culling strategy had the same effect on adult survival and recruitment rates, as a bird killed before the breeding season reduces the breeding population (lower adult survival rate) and prevents the bird from reproducing (lower recruitment rate). When using the breeding culling strategy, recruitment rate was more affected than adult survival rate. This was probably due to immatures being much more susceptible to being shot than adults (Bellrose, 1980; Fox et al., 2014). The compensation hypothesis states that the increased mortality of immatures due to harvesting may be compensated for by a higher natural survival (Cooch et al., 2014), but this hypothesis is not supported by the observations. Overall, there was no evidence of a compensatory mechanism in either context. This could be due to the fact that both populations were far from reaching carrying capacity, meaning that competition for resources was relaxed (Péron, 2013).

With more or less comparable culling pressure, the pre-breeding culling strategy proved to be much more effective. However, this does not mean that the breeding culling strategy was not also effective. Conversely, it proved to be quite efficient in stopping both the population increase and the expansion of the distribution area, and in triggering a significant population decline in a second attempt by slightly increasing the culling pressure. So if winter culling is not possible, which was the case in FR, culling during the breeding season is effective in stopping or reversing the increase in populations of this species. However, the breeding culling strategy is compromised by the highly variable detectability of immatures, as these individuals colonize new ponds. This leads to fluctuations in the recruitment rate and thus jeopardizes the long term efficiency of this strategy. This problem does not exist with the pre-breeding strategy, as the Ruddy ducks form large flocks that occupy a limited number of sites during winter (Johnsgard & Carbonell, 1996).

The response of the two Ruddy duck populations to culling pressure shows that it is necessary to take the culling period into account in order to make correct predictions about demography trajectory. Predicting the effects of harvest pressure on a waterfowl population is then not only a question of the level of harvest, but also the timing of the harvest (Kokko et al., 1998). Unexpectedly, our results suggest that harvesting waterfowl during the breeding season has far less impact on population growth rate than harvesting during winter. If investigations on game birds lead to the same observation, a target for effective management of harvested waterfowl could be an earlier starting and ending of the hunting season.

### 4.3 Implications for waterfowl management

A possible implication of our study concerns the timing of harvest to limit its impact on population growth rates. Unexpectedly, our results indicate that the persistence of waterfowl populations may benefit from earlier openings and closings of the hunting season (see above). However, this conclusion should be taken with caution, as the harvesting process and the resulting pressure are very different between game birds and introduced species. Moreover, an earlier opening would go against the key concept that hunting does not open while young have not yet fledged.

Tracking fluctuations in population abundance is a common tool for determining the conservation status of a population (e.g. Folliot et al., 2022). But tracking abundance alone does not provide enough information to assess the underlying mechanisms behind changes in population growth (Williams et al., 2002). This requires “digging deeper” (Austin et al., 2000), for example, by monitoring individuals to assess parameters such as survival and recruitment rates (e.g. Arnold, 2018). However, monitoring individuals is time-consuming (e.g. Souchay & Schaub, 2016) and not always possible for endangered species. The “apparent sex ratio” method makes it possible to circumvent these disadvantages in dichromatic species with delayed sexual maturity of the males. These species include the White headed duck in southern Spain, which is highly endangered and cannot be disturbed through capture-mark-recapture. Thus, by improving census strategies and disaggregating the effects of variations in adult survival and productivity on population growth using the approach presented here, much more valuable information on the relevance of management strategies could be obtained.

Finally, we would like to emphasize that the main aim of our approach was to decompose population growth into its two main components, and not to provide unbiased estimates of adult survival or recruitment rates. However, if necessary, count surveys could be designed to do this, at least in theory, for species that exhibit observable delayed dichromatism. This trait is very common in dabbling and diving ducks, but the delay in dichromatism is especially large in some species, making the method particularly suited for them, e.g. in stiff-tailed duck, *Oxyura sp.* (Johnsgard & Carbonell, 1996), Tufted duck, *Aythya fuligula*, Black scotter, *Melanitta nigra americana*, Common Goldeneye, *Bucephala clangula americana* (Bellrose, 1980; Johnsgard, 1978). To apply the “apparent sex ratio” method, the time window of censuses must be chosen wisely so that it fits into the period when immature males do not look like adult males yet, which can differ from a species to another. Apart from this precaution, modifying standard monitoring protocols to distinguish between male-like and female-like individuals is almost costless, but worth the effort as it would greatly increase the efficiency of conservation/management actions (Nichols & Williams, 2006).

## Appendix

**Figure 8:**
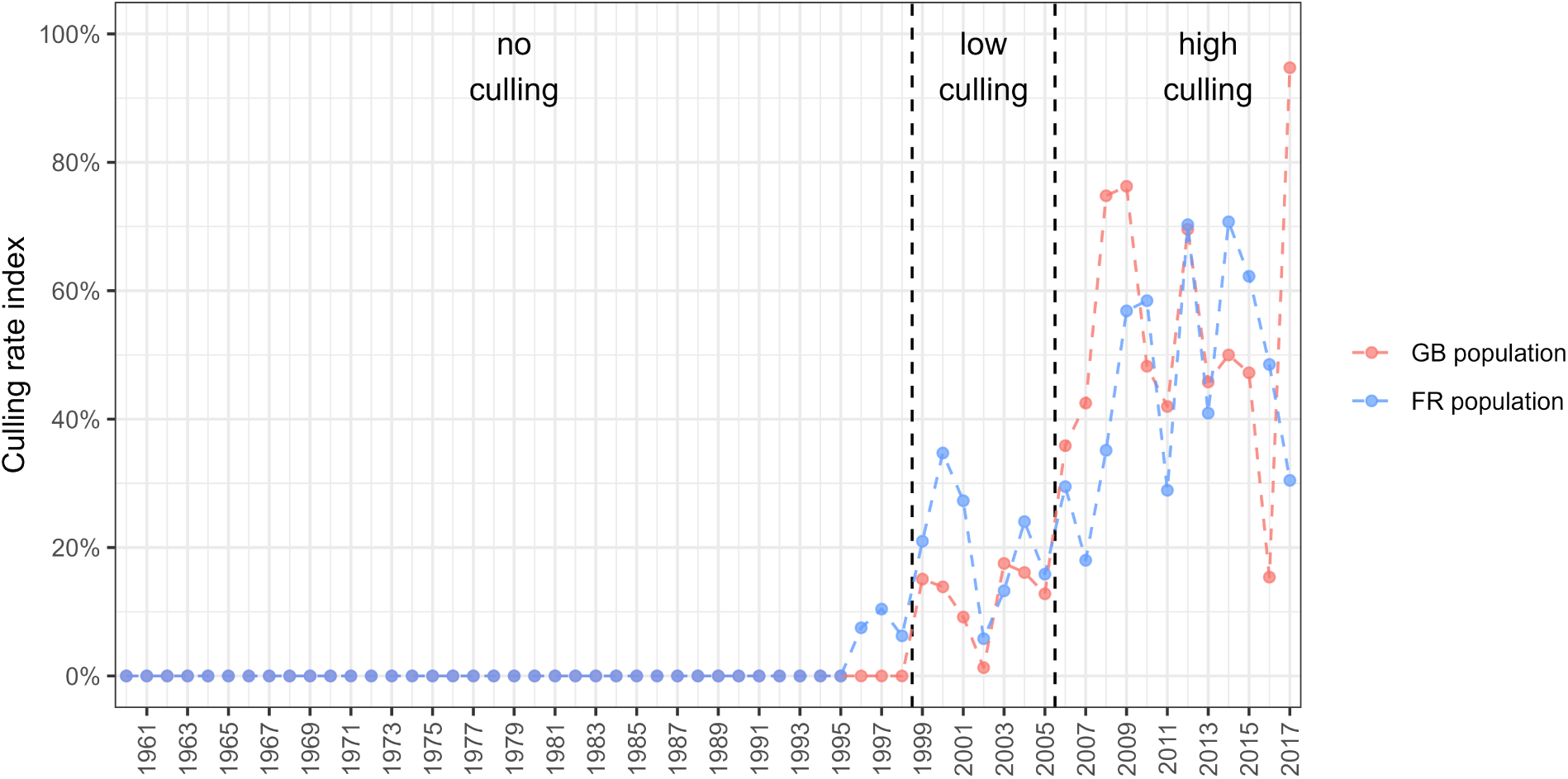
Definition of three periods with different culling pressures using an index based on an estimate of the culling rate on adults (the culling rate on juveniles is not available); as only half of the culled individuals in France were age-determined before 2009, we applied the average age ratio over 2009-2019 to the remaining individuals in order to obtain an estimate of the number of adults in the culling data before 2009; the culling rate increased over time, so we divided the time series into three categories: we defined a “no culling” period before 1999 as the culling rate for both countries was mostly zero and always below 10%, a “low culling” period between 1999 and 2005 as the culling rate for both countries fluctuated around 20%, the culling rate then increased significantly for both countries, therefore we defined a “high culling” period from 2006 onwards

## Acknowledgments

We warmly thank Jay McGowan for his permission to publish his photograph of Ruddy duck. This work was carried out at the suggestion of Jean-François Maillard and Jean-Baptiste Mouronval of the *Office Français de la Biodiversité*, and Jean-Marc Gillier of the *Société Nationale pour la Protection de la Nature*. The authors acknowledge all the contributors of the data collection: in Great Britain, Wildfowl & Wetlands Trust staff past and present, in particular Baz Hughes, Peter Cranswick and Colette Hall, and all the project workers at the Animal and Plant Health Agency and its predecessors & in France, Office Français de la Biodiversité staff past and present, especially Vincent Fontaine, Denis Lacourpaille, Agathe Pirog, Hugo Pichard, Justin Potier, Alexis Laroche, Médéric Lortion, Jules Joly, and Valentin Boniface, as well as the whole team of the Société Nationale pour la Protection de la Nature of the Grand Lieu lake. Thanks to the two anonymous reviewers for their fruitful comments and suggestions.

## Funding

This work was partly funded by the LIFE Oxyura project (LIFE17 NAT/FR/000942) under the LIFE programme.

## Conflict of interest disclosure

The authors declare that they comply with the PCI rule of having no financial conflicts of interest in relation to the content of the article.

## Data, scripts, code, and supplementary information availability

Data and scripts are available online: https://doi.org/10.5281/zenodo.11471723; Tableau et al., 2024

## Notes

https://doi.org/10.5281/zenodo.11471723

### Competing Interest Statement

The authors have declared no competing interest.

### Summary of Updates

insertion of PCI Ecology Badge deletion of line numbering

